# Evidence for *Burkholderia gladioli* pv. *alliicola* Extracellular Detoxification of Thiosulfinates

**DOI:** 10.64898/2026.08.14.744870

**Authors:** Sujan Paudel, Yaritza Franco, Hsiao-Hsuan Jan, Brian Kvitko

**Affiliations:** – Department of Plant Pathology, University of Georgia, Athens. GA, U.S.A.; – Department of Microbiology, University of Georgia, Athens, GA, U.S.A

## Abstract

Onion tissues produce antimicrobial thiosulfinates after tissue damage and cellular decompartmentalization. *Burkholderia gladioli* pv. *alliicola* (Bga), a common onion pathogen, encodes a thiosulfinate tolerance gene (TTG) cluster that protects the bacterium during thiosulfinate exposure. Previous work showed that the TTG cluster contributes to foliar infection but has little effect on infection of onion bulb tissue. To further examine Bga-thiosulfinate interactions in foliar and bulb tissues, we used a thiosulfinate-responsive PaltR-Lux reporter strain to determine when and where Bga encounters thiosulfinates.

In leaves, Bga-induced necrosis was associated with de-repression of the P_altR_-Lux reporter and coincided with a contribution of the TTG cluster to bacterial population size, indicating thiosulfinate exposure during foliar infection. In contrast, TTG mutants and wild-type (WT) strains showed similar growth in scales, and P_altR_-Lux signal declined as scale necrosis progressed, suggesting limited thiosulfinate exposure during bulb colonization. However, when necrosis was induced by the non-native toxin pantaphos, P_altR_-Lux was de-repressed and recovery of the TTG mutant was reduced. These results indicate that Bga encounters thiosulfinates during foliar infection but largely avoids exposure during bulb infection.

Preconditioning the TTG mutant in onion scale tissue did not alter its thiosulfinate *sensitivity in vitro,* arguing against an infection-associated thiosulfinate exclusion mechanism. In contrast, partial rescue of the TTG mutant by the WT strain in zone-of-inhibition co-plating assays suggests extracellular thiosulfinate detoxification. Together, these findings indicate that Bga detoxifies thiosulfinates released during bulb necrosis, limiting thiosulfinate exposure during onion bulb infection. The molecular basis for detoxification and tissue specificity remain unresolved.

## INTRODUCTION

Onion (*Allium cepa* L.), as a widely cultivated vegetable, is crucial to the global agricultural economy. More than a dozen bacterial plant pathogens pose serious threats to the $1 billion onion industry in the U.S.A. (Schwartz et al., 2015; Belo et al., 2023). Since onion bulbs can be stored for extended periods, bacterial diseases may occur under storage conditions, rendering the bulbs unmarketable (Belo et al., 2023). Among the bacterial diseases, slippery skin of onion, caused by the bacterial pathogen *Burkholderia gladioli* pv. *alliicola* (Bga) is commonly observed both in the field and in storage conditions. Bga has also been reported to produce onion foliar symptoms under field and greenhouse conditions (Lee et al., 2005; Paudel et al., 2024a).

Thiosulfinates are reactive organosulfur compounds commonly produced by *Allium* species in response to tissue damage that can be caused by wounding, herbivory, or disease-associated necrosis. These compounds are produced upon tissue damage through a series of chemical and enzymatic reactions leading to distinctive flavors and aromas of onions while also serving as volatile irritants and antimicrobial compounds (Rose et al., 2005; Eady et al., 2008; Leontiev et al., 2018). In onion, cellular damage enables the mixing and reaction of preformed non-volatile precursor cysteine sulfoxide substrates, like isoalliin and propiin, and the carbon-sulfur lyase enzyme alliinase, which are stored in separate sub-cellular compartments to produce thiosulfinates (Lancaster and Collin, 1981; Van Damme et al., 1992; Hughes et al., 2005). Thiosulfinates are volatile antimicrobial compounds that inactivate critical enzymes and deplete the reduced glutathione pool making cells susceptible to oxidative stress (Yamazaki et al., 2010; Müller et al., 2016; Reiter et al., 2020).

Thiosulfinate tolerance gene clusters are a set of thiol redox-associated genes identified in multiple onion-pathogenic bacterial genera. In onion pathogenic *P. ananatis* and multiple *Burkholderia* species, the cluster contributes both to *in vitro* thiosulfinate tolerance and growth in onion juice (Stice et al., 2020; Paudel et al., 2024b). A chemical arms race model has been proposed between the necrotrophic pathogen *P. ananatis* and onion’s thiosulfinate based defense response, wherein thiosulfinates are produced in response to *P. ananatis* induced necrosis. As a counter-defense strategy, *P. ananatis* utilizes the TTG cluster to withstand the toxic thiol stress induced by thiosulfinates (Stice et al., 2020).

Our previous study has shown that TTG clusters are widely distributed among distantly related onion-pathogenic *Burkholderia* species. Surprisingly, and dissimilar to what has been observed in *P. ananatis*, the TTG cluster in all three major onion pathogenic species *B. cepacia*, Bga, and *B. orbicola* were not associated with observable contribution to onion scale colonization (Paudel et al., 2024b). This is unexpected given that the TTG mutants were both impaired in their capacity to grow in onion juice and the Bga TTG genes made a clear contribution to bacterial populations in onion leaves with a lower reported thiosulfinate production capacity than scale and bulb tissue (Cho et al., 2024). This suggests that *Burkholderia* thiosulfinate resistance mechanisms vary based on onion tissue type.

In this study, we hypothesized that Bga may either suppress the production of thiosulfinates in onion scale tissue, inactivate or detoxify thiosulfinates in onion scale tissue, or may exclude the uptake of these compounds into the bacteria. To assess bacterial thiosulfinate exposure, we monitored transcriptional regulation of the thiosulfinate-responsive *altR* promoter region as a *lux* reporter construct. Additionally, pantaphos-containing cell-free spent medium was used as an acute necrosis-inducing factor to examine Bga’s response to non-native onion cell death (Shin et al., 2023). Both *in vitro* Zone of Inhibition (ZOI) and *in planta* foliar coinfection assays were performed to test the potential involvement of Bga in extracellular detoxification of thiosulfinates.

Our findings indicate that Bga is natively exposed to thiosulfinates in onion leaf tissue but not in scale tissue during native Bga-driven necrosis. However, when the non-native necrosis factor pantaphos is used, Bga is exposed to thiosulfinate based on *altR* de-repression, and impaired recovery of the Bga TTG mutant from pantaphos-treated scale tissue. Furthermore, capacity of Bga WT to partially rescue growth of a Bga TTG mutant in ZOI co-plating assays suggests that extracellular detoxificatiuon may play a role in Bga interactions with thiosulfinates.

## RESULTS

### Thiosulfinate exclusion and detoxification models for investigating Bga exposure to thiosulfinates during native onion necrosis

We proposed the thiosulfinate exclusion model and thiosulfinate detoxification model as two hypotheses that could explain why Bga in scale tissue are not exposed to thiosulfinates under native necrosis (Fig. 1). In the exclusion model, Bga colonizing scale tissue develop mechanisms, for instance, formation of a biofilm, which physically excludes thiosulfinates from entering the Bga cells. Alternatively, the thiosulfinate detoxification model posits that Bga secretes factors that either detoxify thiosulfinates or prevent thiosulfinate production in necrotic tissue, thereby neutralizing their toxic effects before they reach the cell.

**Fig. 1:**
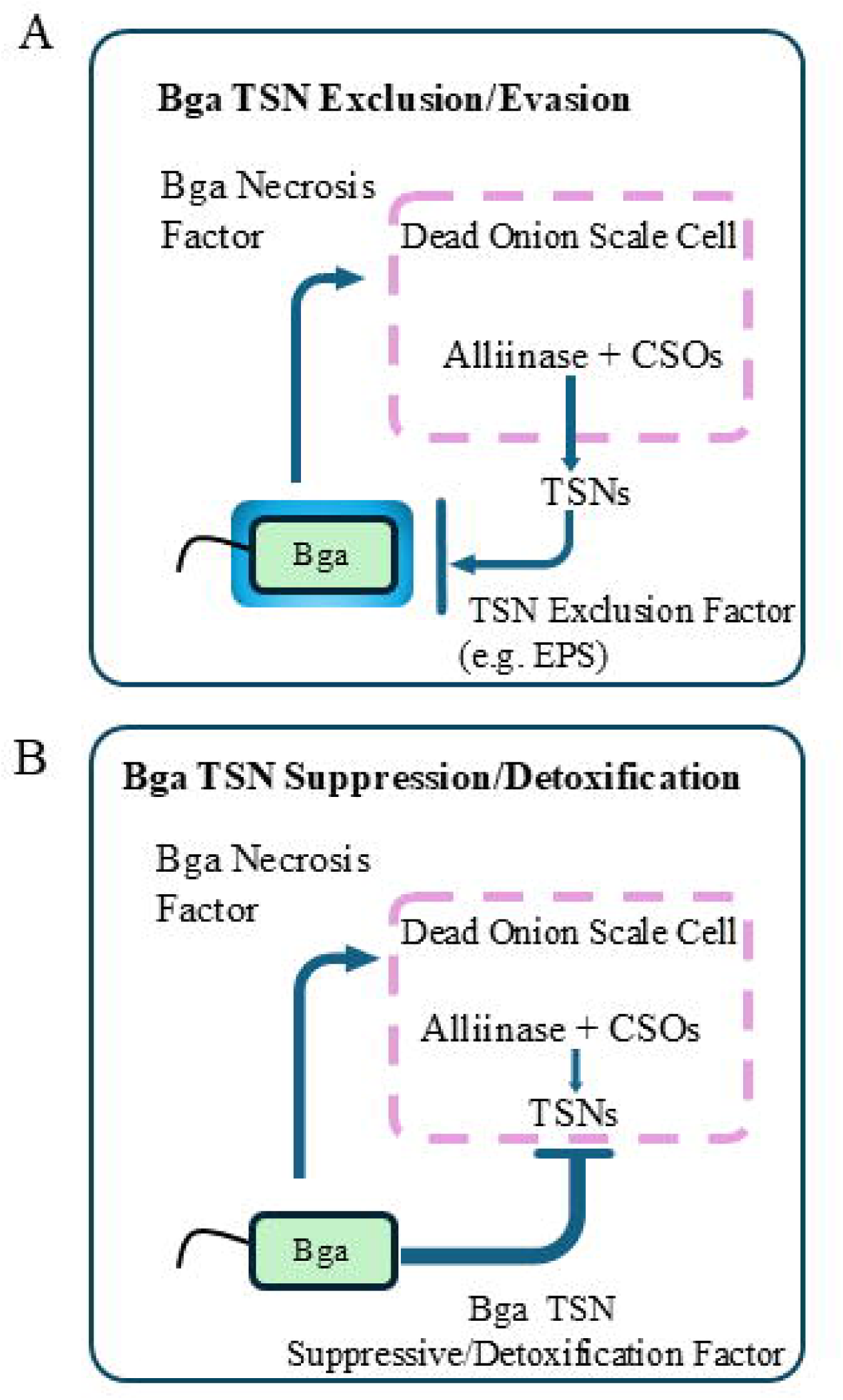
Proposed models for thiosulfinate resistance mechanisms of *Burkholderia gladioli* pv. *alliicola* (Bga) in necrotized onion scales tissue. **A**, In the Bga thiosulfinates (TSNs) exclusion model, the bacterium evades or excludes thiosulfinates exposure using the TSNs exclusion factor such as exopolysaccharides (EPS) and biofilm formation. **B**, In the Bga TSN suppression/detoxification model, the bacterium inhibits the TSN precursor CS lyase enzyme alliinase or detoxify/inactivate thiosulfinates in scale tissue.

### De-repression of Bga P*_altR_*^PNA 97-1R^ Lux construct was observed with onset and progression of necrosis in onion seedling assays

Jan et al. 2025 established that the P*_altR_*^PNA 97-1R^Lux reporter is repressed by the AltR transcription factor and derepressed upon exposure to thiosulfinates. We generated a similar reporter construct based on the P*_altR_*^Bga20GA0385^ native promoter, which showed similar behavior but a lower luminescence signal than the P*_altR_*^PNA 97-1R^Lux reporter. While both *P. ananatis* and Bga P*_altR_* promoters were responsive, P*_altR_*^PNA 97-1R^ was selected for its superior signal with minimal background (Fig. S1). As the Bga TTG cluster contributed to *in planta* bacterial population in onion leaves, we hypothesized that upon onset and progression of foliar necrosis, Bga would be exposed to thiosulfinates and the 20GA0385 P*_altR_*^PNA 97-1R^Lux construct would be de-repressed. Seedlings aged 8-12 weeks were either inoculated with 20GA0385 P*_altR_*^PNA 97-1R^Lux construct to monitor the thiosulfinate exposure or a constitutively bioluminescent 20GA0385 P*_ompA_* Lux strain to monitor the bacterial colonization (P*_ompA_* Lux) signal from day 1 to day 3 post-inoculation. With the onset of necrosis 24 h post inoculation, de-repression of the P*_altR_*Lux reporter construct was observed in 80% of the inoculated seedlings. Along with the progression of necrosis, de-repression was also consistently observed. By day 3, as the inoculated seedlings were severely necrotic and wilting, P*_altR_* de-repression was observed in 69% of the inoculated seedlings leaves (Fig. 2A). Bioluminescence was also consistently observed in the seedlings inoculated with the P*_ompA_* constitutive reporter construct across all three days post inoculation (Fig. 2B).

**Fig. 2:**
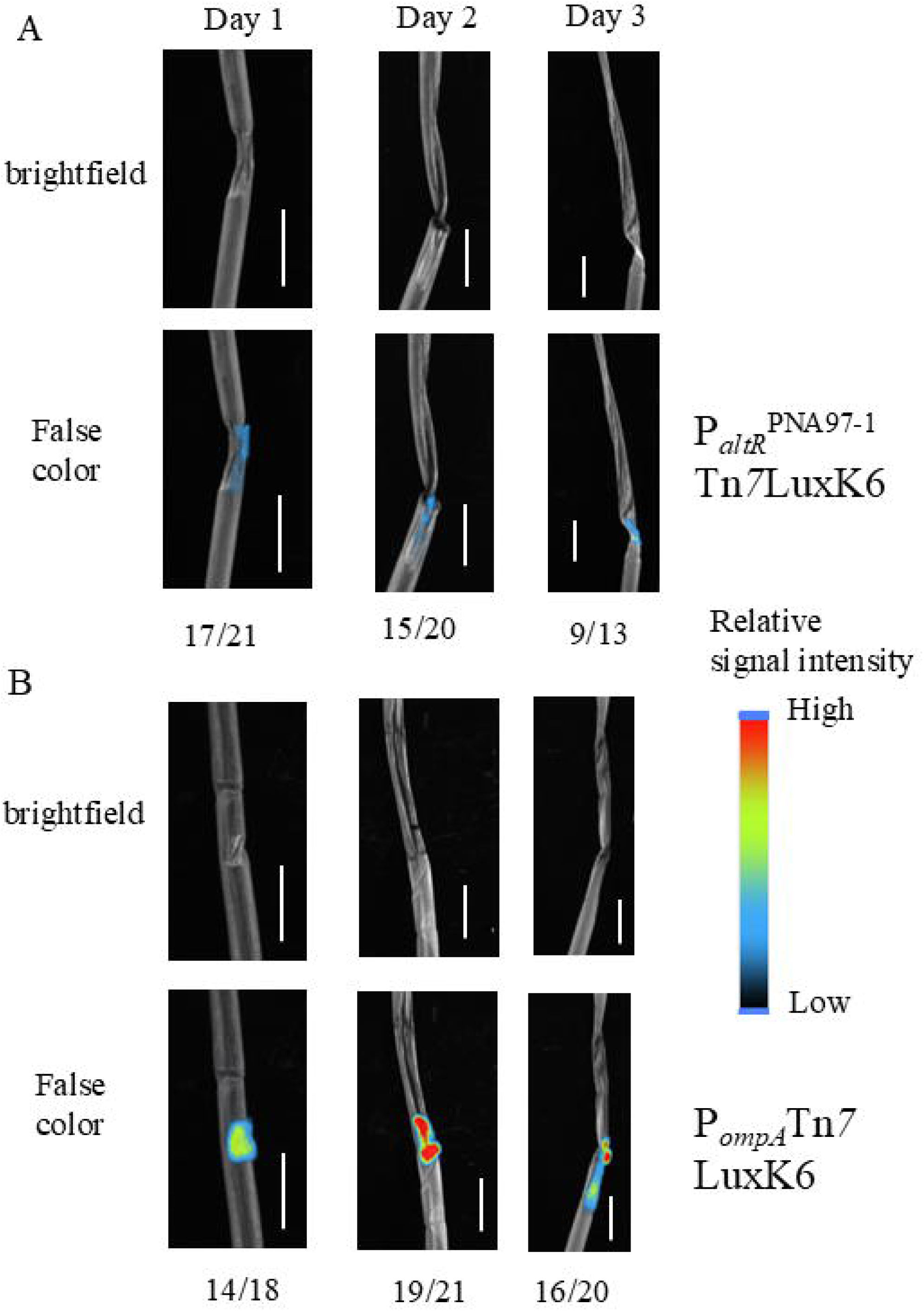
De-repression of Bga *altR* promoter reporter construct is consistent with the initiation and progression of foliar necrosis. Representative brightfield and corresponding false color image of seedlings inoculated with **A** 20GA0385 P*_altR_*^PNA 97-1R^Tn*7*LuxK6 strain and **B** 20GA0385 P*_ompA_*Tn*7* LuxK4 strain from day 1 to day 3 post inoculation. The number in the numerator is the number of inoculated leaves that showed positive bioluminescence signal for samples visualized on corresponding day. The number in the denominator is the total number of inoculated leaves. Legend bar represents the relative intensity of visible signal from low to high. Scale: 1 cm. Bioluminescence signal visualized using Newton 7.0 camera and analyzed using Kuant v 2.5 software.

### De-repression of Bga P*_altR_*^PNA 97-1R^ displayed different patterns during Bga native necrosis than during pantaphos cell free spent medium induced necrosis in scale tissue

Pantaphos cell free spent medium was used as a non-native necrosis inducing factor in the red scale necrosis assay to monitor the de-repression of 20GA0385 P*_altR_*^PNA 97-1R^Lux promoter reporter in the necrotized onion scale tissue. Scales inoculated with the mixture of pantaphos containing cell free spent medium and 20GA0385 P*_altR_*^PNA 97-1R^ de-repression was observed in all the inoculated scales commensurate with the onset of pantaphos-induced scale necrosis 48 h post inoculation. The number of inoculated scales that showed positive bioluminescence signal was reduced to 6 of 18 at 72 h post inoculation (Fig. 3A). The constitutive bioluminescent construct 20GA0385 P*_ompA_* Lux showed bioluminescence signal in all the inoculated scales at all time points post inoculation (Fig. 3B).

**Fig. 3:**
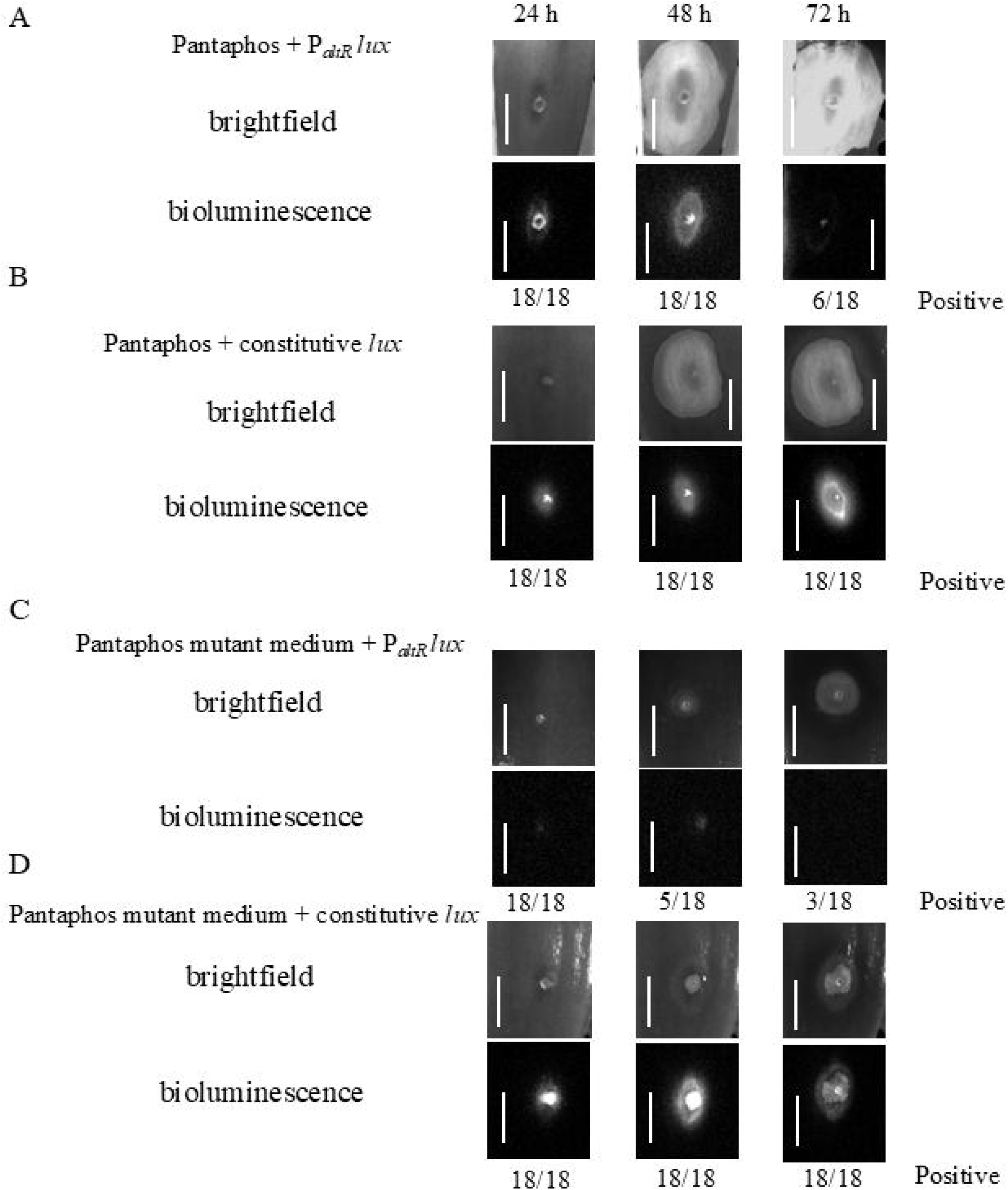
The de-repression pattern of Bga *altR* promoter reporter construct is distinct during pantaphos induced necrosis as compared to native scale necrosis. Representative brightfield and corresponding bioluminescence image of scales inoculated with A, Cell free spent medium of PNA 97-1R P_SNARE_ *hvr* and Bga P*_altR_*^PNA 97-1R^Tn*7*LuxK6; B, Cell free spent medium of PNA 97-1R P_SNARE_ *hvr* and Bga P*_ompA_*Tn*7*LuxK4; **C,** Cell free spent medium of PNA 97-1R Δ*hvrE* P_SNARE_ *hvr* and Bga P*_altR_*^PNA 97-1R^Tn*7*LuxK6 and **D,** Cell free spent medium of PNA 97-1R Δ*hvrE* P_SNARE_ Δ*hvrE* strain and 20GA0385 P*_ompA_*Tn*7* LuxK4. The number in the numerator is the number of inoculated scales that showed a positive bioluminescence signal for samples visualized on the corresponding day. Brightness and contrast of the presented scales were adjusted to enhance visibility for presentation. The number in the denominator is the total number of inoculated scales across three independent experimental repeats. For each panel, the brightfield and bioluminescence series each track a single representative scale continuously from 24 to 72 hours post-inoculation. Samples were imaged under 2 min exposure for the 24 h time point and under 5 min exposure for the rest of the time points. Scale: 1 cm.

Cell-free spent medium prepared from of *P. ananatis* Δ*hvrE* mutant strain defective in pantaphos production (Shin et al., 2023) was mixed with Bga *altR* promoter reporter strain and inoculated into scale tissue. This spent medium provides a pantaphos-negative control for monitoring Bga responses during native necrosis. Minor de-repression was observed 24 h post-inoculation in all the inoculated scales. After 24 h, the de-repression signal decreased daily and was largely lost by day 3 (Fig. 3C). The bacterium was able to successfully colonize onion scales in the presence of P_SNARE_ Δ*hvrE* cell-free spent medium, as evident from the bioluminescence signal observed in all the inoculated scales across all time point (Fig. 3D).

### The recovery of the Bga TTG mutant from scale tissue was reduced compared to WT strain during Pantaphos-induced necrosis

Red scale assay with the cell-free spent medium was performed to test the recovery of WT and TTG mutant population from the pantaphos-necrotized onion scale tissue. The recovery of the Bga TTG mutant treated with pantaphos spent medium mixture was significantly lower compared to the WT strain at day 2 and day 3 post-inoculation (Fig. 4A). However, when the cell free spent medium of the Δ*hvrE* mutant was used, the recovery of the TTG mutant from inoculated scale tissue was similar to the WT strain from day 1 to day 3 post-inoculation (Fig. 4B).

**Fig. 4:**
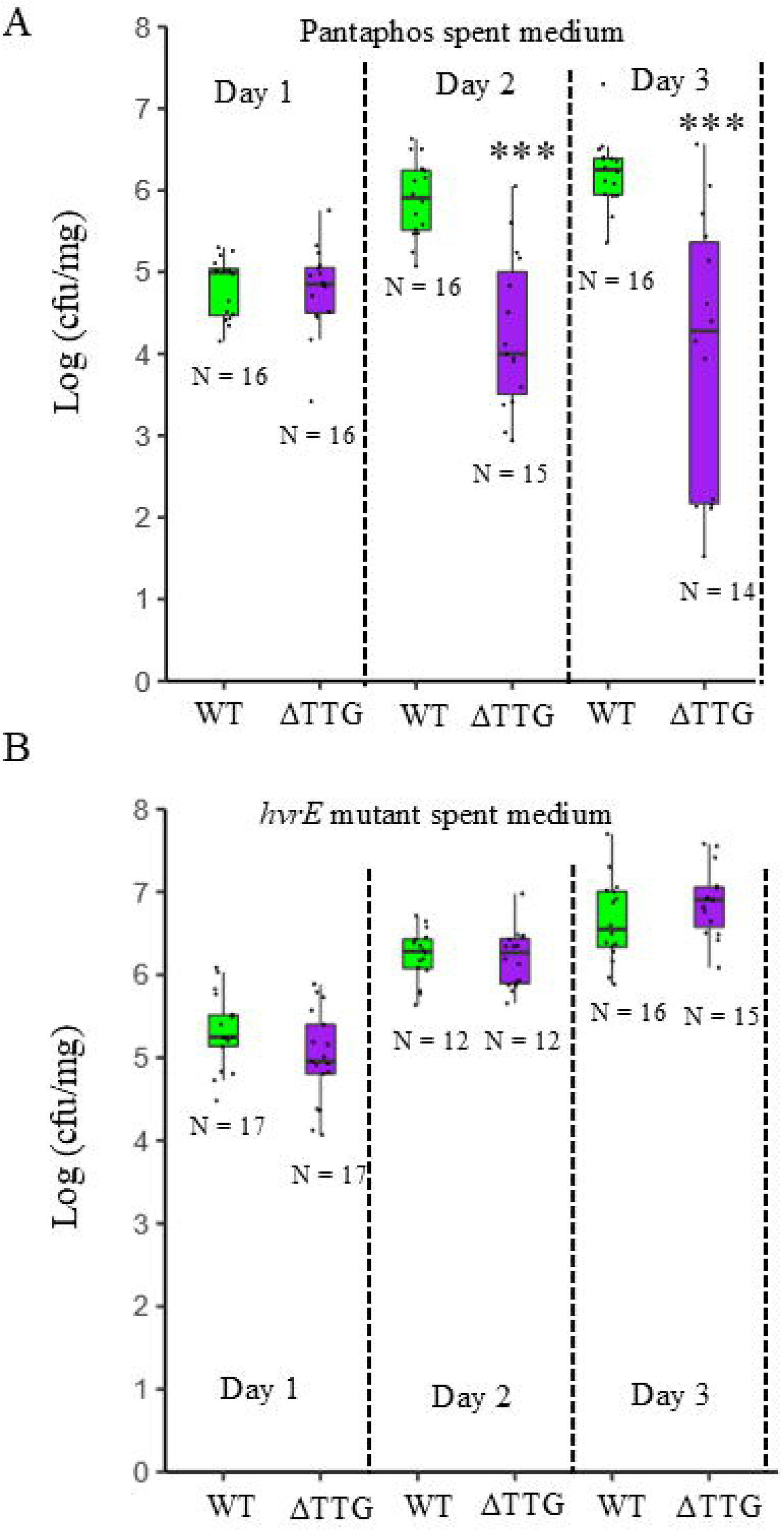
The TTG mutant exhibits impaired scale recovery in the presence of pantaphos cell-free spent medium, which serves as an external necrosis-inducing factor. Bacterial population recovery of Bga WT Tn7 LuxK4 and ΔTTG Tn7 LuxK4 strains from inoculated scales was monitored from day 1 to day 3 post-inoculation containing cell-free spent medium from **A**, PNA 97-1R P_SNARE_ *hvr* and **B**, PNA 97-1R Δ*hvrE* P_SNARE_ *hvr*. N is the total number of scales inoculated across four independent experimental repeats. Level of significance: 0 ‘***’ 0.001 ‘**’0.01 ‘*’ 0.05.

### Preconditioning the Bga TTG mutant in scale tissue did not impact its growth behavior in onion juice growth assays

To determine if preconditioning the Bga TTG mutant in the scale tissue would confer protection from thiosulfinate exposure via exclusion, Bga TTG mutant bacteria were recovered directly from the symptomatic area of inoculated scale tissue and then exposed to onion juice. The TTG strain preconditioned in the scale tissue for 3 days showed dramatically reduced growth relative to the similarly preconditioned WT strain. The growth pattern of the non-conditioned WT and TTG mutant strains in the scale tissue resembled the previously observed pattern for WT and TTG strains (Fig. 5) (Paudel et al., 2024b).

**Fig. 5:**
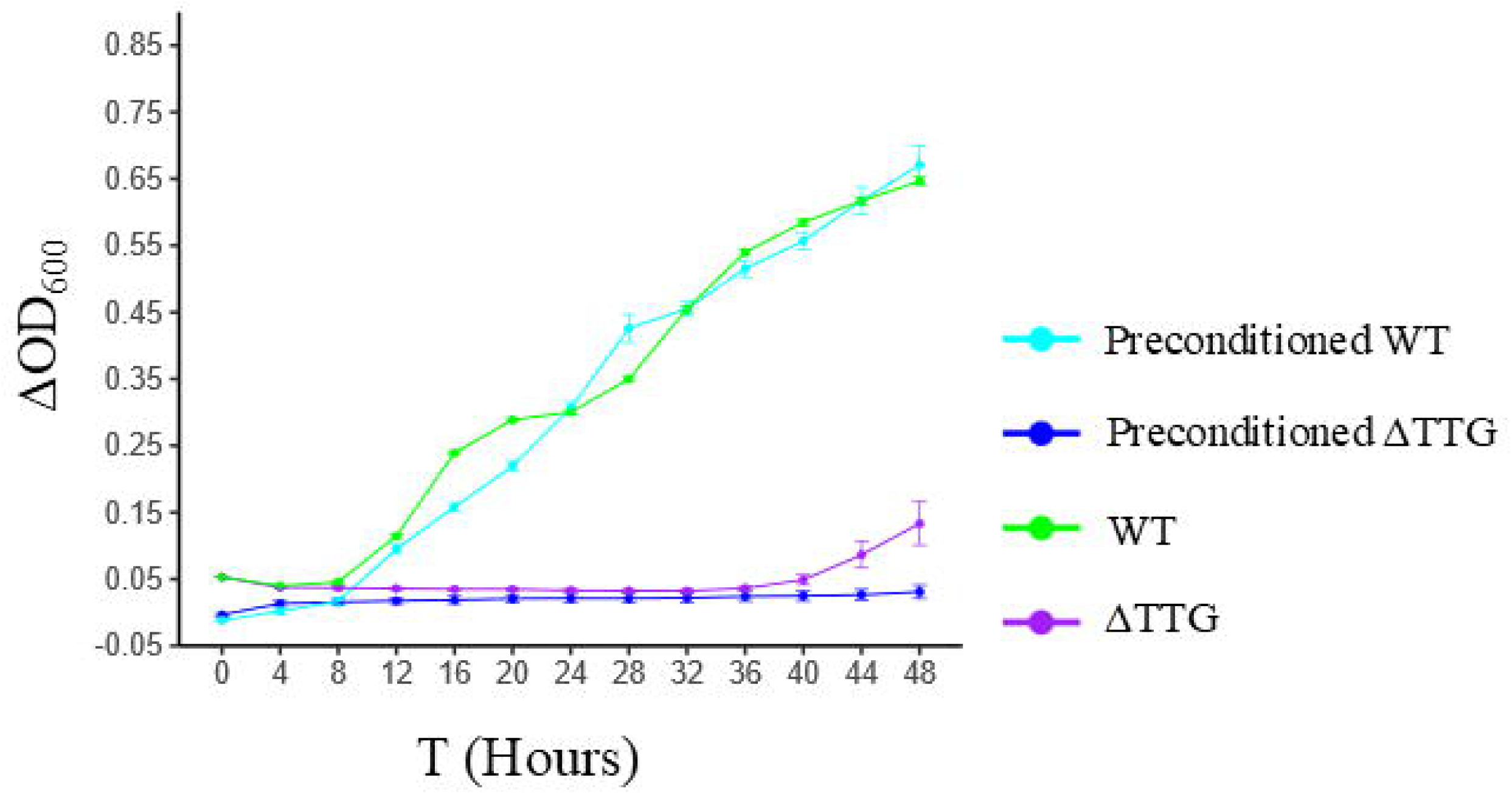
Preconditioning the TTG mutant in the scales didn’t change its growth behavior in onion juice. Line graph showing the growth of preconditioned/non-conditioned Bga WT and TTG mutant strains in onion juice over the period of 48 hours. Absorbance values were normalized by subtracting the average absorbance of the kanamycin-treated normalization control or half-strength onion juice (negative control) from each well. The graph presents the average values from two independent experimental repeats at each time point. Error bars represent ± standard error. Cyan represents preconditioned WT strain, blue represents preconditioned TTG mutant strain, green represents non-conditioned WT strain and violet represent non conditioned TTG mutant strain.

### The Bga WT strain was able to partially rescue the TTG mutant in thiosulfinate *in vitro* zone of inhibition co-plating assays

A ZOI co-inoculation assay was conducted to evaluate the thiosulfinate detoxification model (Fig. 1B). We reasoned that if the Bga WT strain secretes factors that facilitate extracellular detoxification of allicin, the presence of the WT strain would confer cross-protection to the allicin-sensitive TTG mutant, thereby reducing the ZOI area of the co-inoculation mix observed *in vitro.* In the ZOI co-plating assay, the ZOI area of the TTG mutant from the WT/TTG co-inoculation mix was significantly lower than the ZOI area of the TTG mutant-only strain. The ZOI area of TTG strain in the co-plating mix was still significantly larger than the WT-only strain, suggesting the rescue phenotype is only partial (Fig. 6).

**Fig. 6:**
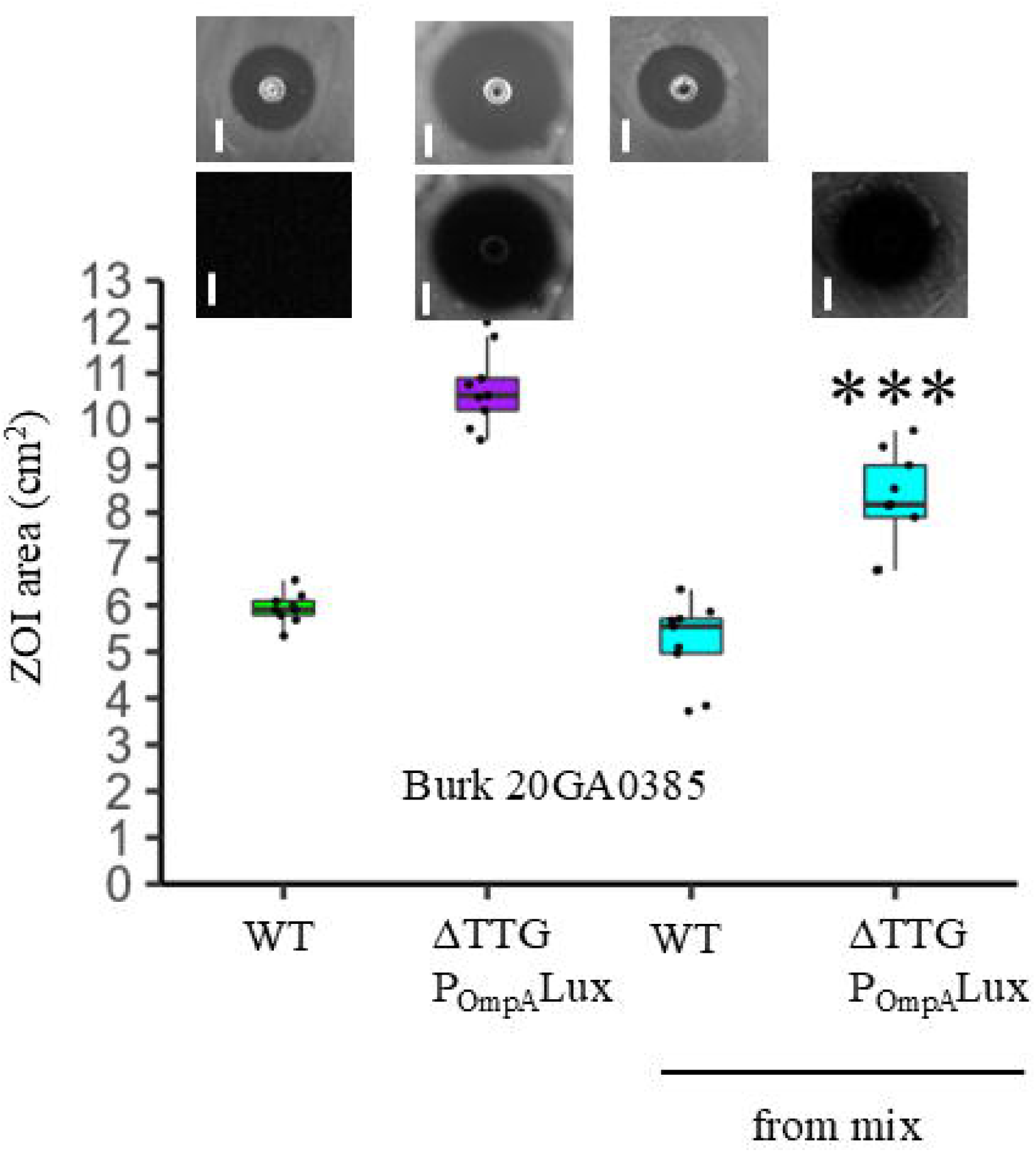
Bga WT strain partially rescues the phenotype of TTG mutant strain in the zone of inhibition (ZOI) co-inoculation assay. Box plot showing the allicin ZOI area of Bga 20GA0385 WT, TTG mutant, WT from co-inoculation mix and TTG mutant from co-inoculation mix. Representative brightfield and/or bioluminescence images of ZOI plates treated with corresponding strains are presented above the box. Bioluminescence images were taken under 2 minutes of exposure. Experimental data from three independent repeats (n = 9) are presented. Statistical differences in ZOI area between the TTG mutant and the TTG mutant in the co-inoculation mix were calculated using the pairwise t-test function in Rstudio. Level of significance: *** < 0.001. Scale: 1 cm.

### Bga WT strain does not restore TTG mutant populations in onion foliar co-inoculation assays

We performed leaf co-inoculation assays with onion seedlings and mature onion plants to assess whether the Bga WT strain could rescue the Bga TTG mutant from thiosulfinate exposure and partially restore its populations in tissue. The CFU/cm recovery of TTG mutant from the WT and TTG co-inoculation mix was similar to the recovery of TTG treatment alone. As expected, the bacterial population of the Bga WT strain from the infected seedling sample was significantly higher than the Bga TTG mutant (Fig. 7). The TTG mutant population was also not rescued by the WT strain in the mature onion leaves co-inoculation assay (Fig. S2)

**Fig. 7:**
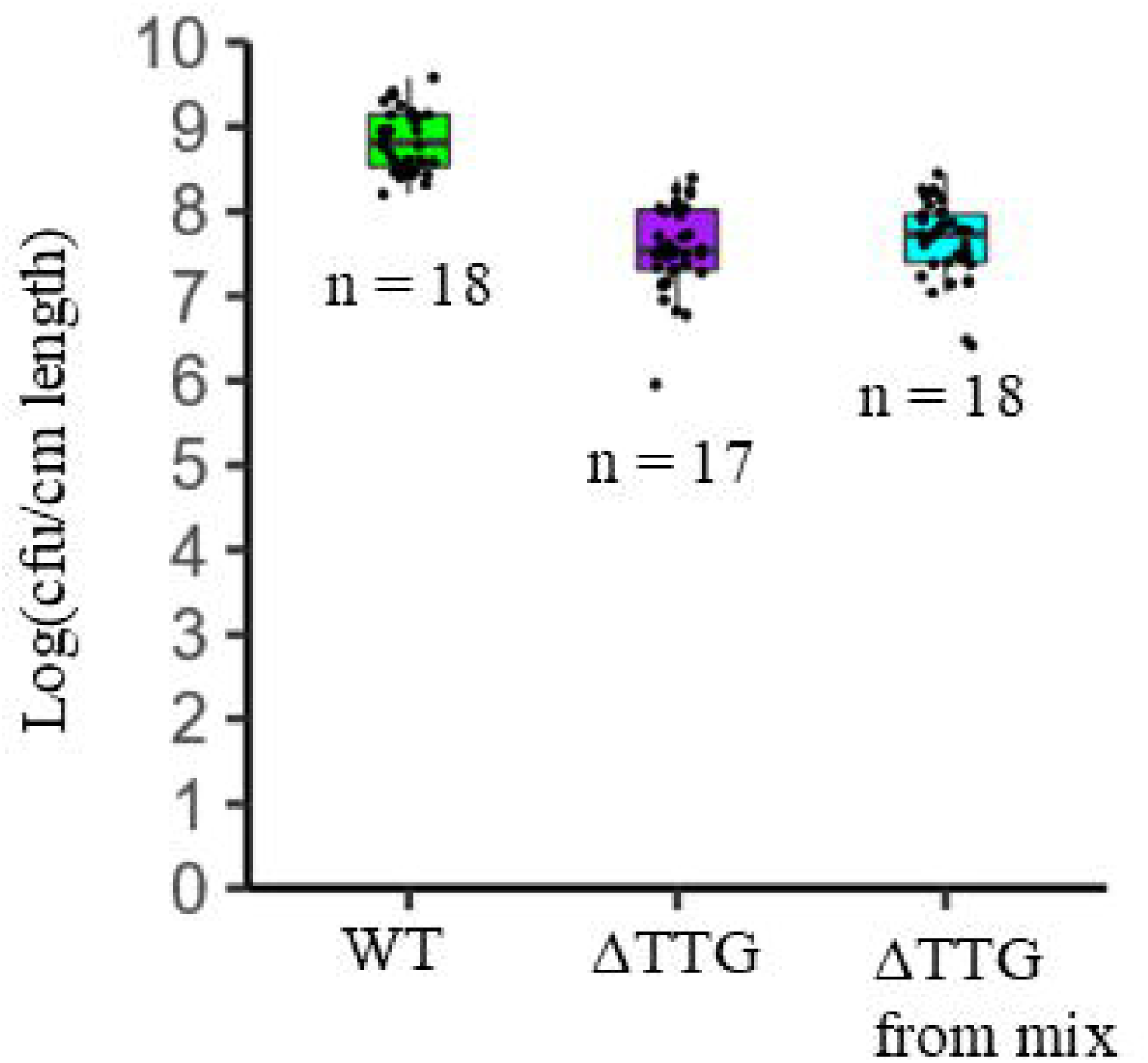
Bga WT strain didn’t rescue the population of TTG mutant from the infected seedlings in the co-infection assay. Box plot showing the CFU/cm of seedling length recovery of Bga WT, TTG mutant, and TTG mutant from co-infection mix after three days post-inoculation. N is the total number of observations across three independent experimental repeats.

## DISCUSSION

The Bga TTG cluster contributes to bacterial populations and development of necrosis symptoms in onion leaves but not in onion scales. This is surprising as the thiosulfinates production potential in the bulb tissue has been reported to be is much higher compared than leaf tissue (Cho et al., 2024). The stark contrast in the requirement for the TTG cluster between leaf and scale tissue suggests a fundamental difference in the way Bga interacts with thiosulfinates in these two environments. We proposed the thiosulfinate exclusion model or thiosulfinate detoxification model as potential explanations (Fig. 1). We used the *altR* Lux promoter reporter constructs, pantaphos cell-free spent medium as a non-native necrosis inducing factor, and ZOI co-plating assays, to infer potential mechanisms for the tissue specific virulence roles of the Bga TTG cluster. Under native necrosis, de-repression of Bga *altR* promoter reporter construct in the scale tissue was weak and transient dissipating after the appearance of Bga-induced native necrosis symptoms. However, in the presence of pantaphos as the onion scale necrosis inducing factor, the de-repression of Bga *altR* reporter was pronounced with the onset of pantaphos-induced necrosis symptoms. In addition, under pantaphos-induced necrosis, the recovery of the TTG mutant was significantly lower than that of the WT strain. Incubating the TTG strains in onion scales prior to exposing them to onion juice did not rescue their growth impairment. This all suggests that an adaptive Bga thiosulfinate exclusion mechanism is unlikely to be at play in onion scales. In the ZOI co-inoculation assay, the Bga WT strain was able to partially rescue the thiosulfinate sensitivity phenotype of TTG mutant strain suggesting the potential extracellular detoxification of thiosulfinates by Bga WT strain.

The *P. ananatis altR* promoter reporter was utilized to determine if the bacteria were exposed to thiosulfinates released from the onion scale tissue upon Bga infection. In the presence of thiosulfinates, the *altR* promoter reporter construct was de-repressed as seen by the bright bioluminescent ring in the edge of ZOI area (Fig. S1). Consistent with the positive contribution of the TTG cluster to Bga virulence in leaf tissue, de-repression of P*altR* was observed in leaf tissue from day 1 to day 3 post inoculation (Fig. 2A).

Following the confirmation of Bga thiosulfinate exposure in foliar tissue, we investigated if similar interactions occur during Bga scale necrosis. We compared the de-repression of Bga P*_altR_* reporter during scale necrosis induced by external necrosis factor, pantaphos against cell free spent medium of *hvrE* mutant as a pantaphos free control (Shin et al., 2023). We observed a strong bioluminescence signal of the Bga *altR* promoter reporter construct with pantaphos induced necrosis suggesting the bacterium is exposed to thiosulfinates (Fig. 3A). This is contrary to what was observed during Bga induced native necrosis where the de-repression of *altR* reported construct diminished with onset and progression of scale necrosis (Fig. 3C). We also found that the recovery of TTG mutant was significantly reduced compared to the WT strain during pantaphos-induced necrosis (Fig. 4). The contrasting patterns of de-repression and differential Bga recovery observed during native scale necrosis vs. pantaphos-induced necrosis suggest that Bga exclusion of thiosulfinates in scale tissue is unlikely.

To further investigate whether the bacterium physically excludes the thiosulfinate exposure through induced tolerance or biofilm-mediated protection, we performed a scale pre-conditioning assay. If the bacterium develops protective mechanisms against thiosulfinate exposure in the scale tissue, the growth of a scale pre-conditioned TTG mutant would be expected to remain improved in onion juice. However, the growth of TTG mutant, when sampled from inoculated scale tissue and exposed to half strength onion juice, was dramatically reduced compared to the WT strain. The growth pattern observed for the pre-conditioned TTG mutant was similar to TTG mutant sampled from culture media plate (Fig. 5) (Paudel et al., 2024b). While mechanical homogenization of scale tissue could potentially disrupt protective structures like biofilms, the de-repression of *altR* reporter in the relatively intact scale tissue during pantaphos induced necrosis reinforces the interpretation that Bga thiosulfinate exclusion in the scale tissue is unlikely. The findings from preconditioning and pantaphos scale assay were inconsistent with an adaptive thiosulfinate exclusion model. We therefore proposed a second hypothesis, the Bga thiosulfinate detoxification model, in which Bga potentially suppresses thiosulfinate production or detoxifies thiosulfinates or their precursors within the scale tissue. When we utilized genetically engineered TTG mutant for the ZOI co-plating assay, we saw that Bga WT strain partially rescues the ZOI thiosulfinate sensitivity phenotype of TTG mutant strain (Fig. 6). This observation suggests that the Bga WT strain may contribute extracellularly to the detoxification of allicin present in the ZOI assay, potentially through secreted factors or enzymatic activity. It is also important to note that the ZOI *in vitro* assay does not contain the enzyme alliinase which suggest the rescue mechanism is independent of suppression or inhibition of alliinase. The recovery of the Bga TTG mutant from the WT and TTG co-inoculation mix in seedling leaf tissue was similar to the recovery of TTG mutant alone (Fig. 7). The rescue effect of the Bga WT strain was also not observed in the mature onion leaves co-inoculation assay (Fig. S2). The lack of rescue effect may be explained by the difference in composition of the agar media and apoplastic environment. While the agar plate offers relatively unobstructed medium for the diffusion of potential detoxification factor, the plant apoplast may limit the mobility of these factors.

Our study delves deeper into the mechanisms behind Bga’s tissue-specific virulence, we use bioluminescence to “shed light” on its interactions with onion thiosulfinates. Using *altR* promoter reporter and pantaphos cell free spent medium, we observed that Bga *altR* promoter de-repression was enhanced in pantaphos necrotized scale tissue, while the TTG mutant was more susceptible to thiosulfinate exposure, suggesting that Bga does not rely on exclusion but may inactivate thiosulfinates released natively within scale tissue. This hypothesis is further supported by the observation that preconditioning TTG mutants in scale tissue did not improve their survival upon exposure to thiosulfinate in onion juice, implying that Bga’s resistance mechanism is not impacted by environmental adaptation. In contrast to the slower, gradual symptom development observed during Bga-induced native scale necrosis, pantaphos-induced necrosis progresses rapidly and extensively, suggesting that the resulting thiosulfinate exposure may surpass the threshold that Bga can intrinsically tolerate. It is possible that Bga secretes enzymatic or chemical compounds capable of inactivating or detoxifying the produced thiosulfinates, or that it prevents their formation within onion scales. Future experiments could focus on screening for the Bga secreted factors that could inactivate or detoxify thiosulfinates. Furthermore, investigating the bacterial transcriptional response using RNA-seq could provide insight into differences in onion tissue-specific gene regulation.

## MATERIALS AND METHODS

### Bacterial growth conditions

Bacterial strains and plasmids utilized in the study are listed in Table 1. Primers and synthesized double-stranded dsDNA fragments are listed in Table 2. *E. coli* strains MaH1, RHO5, all *Burkholderia* and *Pantoea* strains used in the study were grown in Luria-Bertani (LB; per liter, 10 g of tryptone, 5 g of yeast extract and 5 g of NaCl, 15 g agar) or LM (per liter, 10 g of tryptone, 6 g of yeast extract, 1.193 g of KH_2_PO_4_, 0.6 g of NaCl, 0.4 g of MgSO_4_.7H_2_0, and 18 g of agar) media. *E. coli* strains were grown at 37^0^C and *Burkholderia* and *Pantoea* strains were grown at 30^0^C. Antibiotics and chemicals were supplemented with the media using the following final concentrations per milliliter: 50 µg of kanamycin, 10 µg of gentamicin, 100 µg of ampicillin, 25 µg of trimethoprim, and 100 to 200 µg of diaminopimelic acid (DAP) and 40 to 60 µg of rifampicin, as appropriate.

**Table 1:** Strains and plasmids used in the study.

| *E.coli* strain | Purpose | Reference |
| --- | --- | --- |
| MahI | DH5α | (Kvitko et al., 2012) |
| RHO5 | *pir116*, DAP-dependent conjugation strain | (Kvitko et al., 2012) |
| RHO3pTNS3 | DAP dependent conjugation strain SM10<br>derivative with Tn7 transposase helper<br>plasmid (Amp<sup>R</sup>) | (López et al., 2009) |
| 20GA0385 | *Burkholderia gladioli* pv. *alliicola* (Bga) WT strain (Rf<sup>R</sup>) | (Paudel et al., 2024b) |
| 20GA0385<br>ΔTTG | TTG mutant derivative of Bga 20GA0385 (Rf<sup>R</sup>) | (Paudel et al., 2024b) |
| PNA 97-1R | *Pantoea ananatis*, WT strain (Rf<sup>R</sup>) | (Stice et al., 2020) |
| 20GA0385<br>P<sub>ompA</sub>Tn7Lux<br>K6 | *ompA* promoter driven constitutive bioluminescent Bga<br>20GA0385 strain (Rf<sup>R</sup>, Km<sup>R</sup>) | This study |
| 20GA0385<br>ΔTTG<br>P<sub>ompA</sub>Tn7Lux<br>K6 | *ompA* promoter driven constitutive bioluminescent Bga<br>20GA0385 ΔTTG strain (Rf<sup>R</sup>, Km<sup>R</sup>) | This study and<br>(Bruckbauer et al.,<br>2015) |
| 20GA0385<br>P<sub>altR</sub><sup>PNA 97-1R</sup><br>Tn7LuxK6 | *Burkholderia altR* promoter reporter construct with *P. ananatis* PNA 97-1R *altR* promoter region (Rf<sup>R</sup>, Km<sup>R</sup>) | This study |
| 20GA0385<br>P<sub>altR</sub><sup>20GA0385</sup><br>Tn7LuxK6 | *Burkholderia altR* promoter reporter construct with Bga<br>20GA0385 *altR* promoter region | This study |
| PNA 97-1R<br>::P<sub>SNARE</sub> *hvr* | Hivir IPTG inducible *P. ananatis* strain with pSNARE<br>inserted in front of *hvrA* gene of pantaphos biosynthesis<br>gene cluster (Rf<sup>R</sup>, Tmp<sup>R</sup>). Used to generate cell free spent<br>medium for co-infection assays in this study. | (Shin et al., 2023) |
| PNA 97-1R<br>Δ*hvrE* ::<br>P<sub>SNARE</sub> *hvr* | Hivir IPTG inducible *P. ananatis* Δ*hvrE* mutant strain with<br>P<sub>SNARE</sub> inserted in front of *hvrA* gene of pantaphos<br>biosynthesis gene cluster (Rf<sup>R</sup>, Tmp<sup>R</sup>). Used to generate cell<br>free spent medium for co-infection assays in this study. | (Shin et al., 2023) |
| pTn5.7LuxK<br>6 | Promoter capture vector, R6K replicon (Km<sup>R</sup>) | KC332287.1 |

**Table 2.**
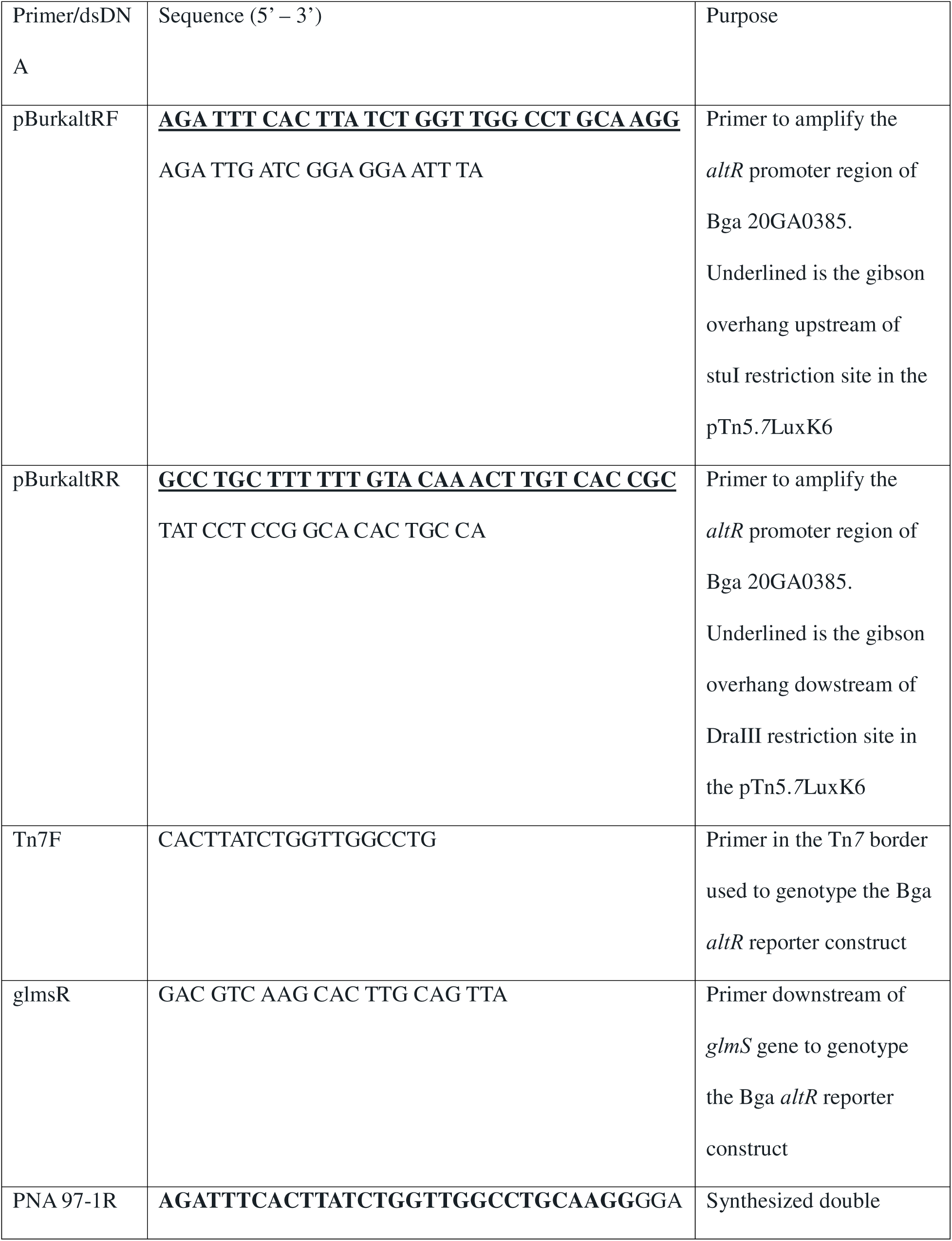
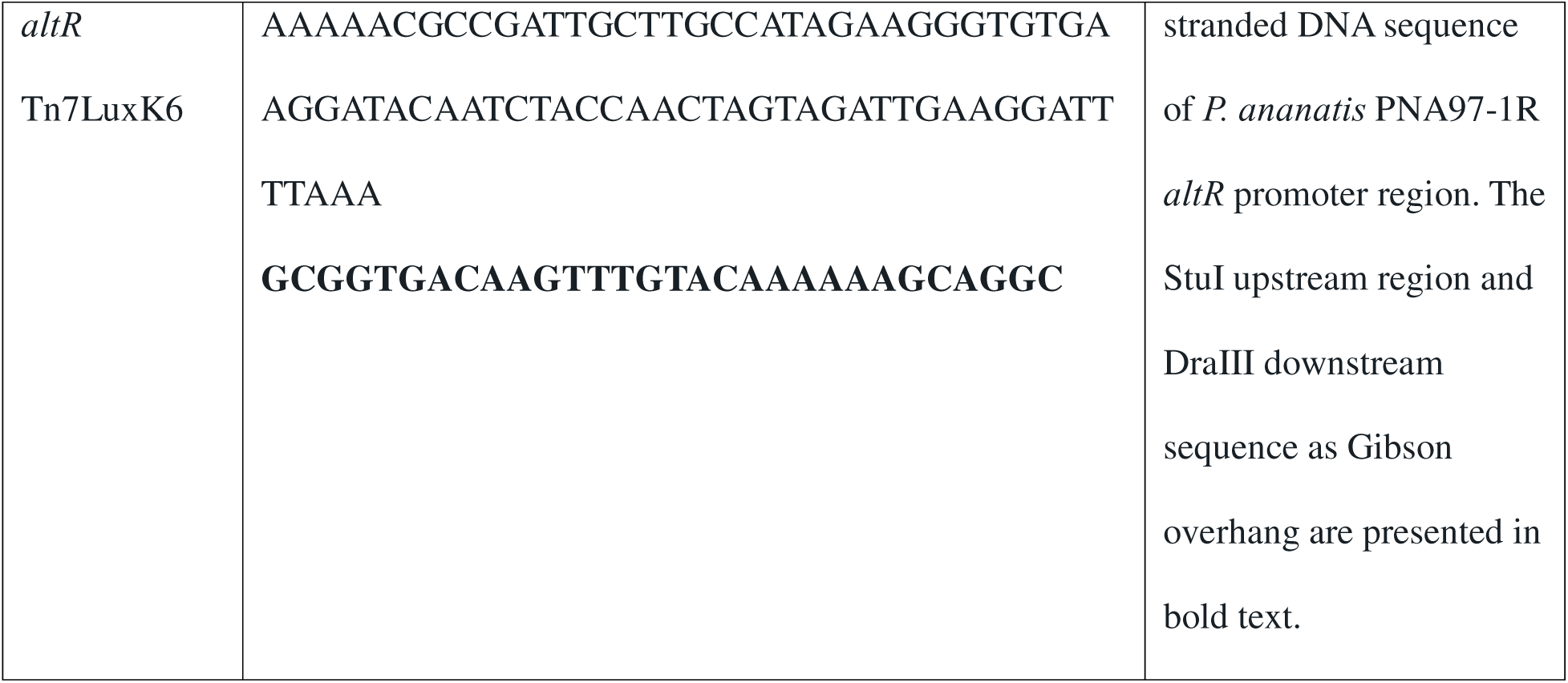
Primers and double stranded synthesized DNA used in the study.

### Plant growth conditions

Onion seedlings for leaf infection and co-inoculation assays were grown and maintained following procedures described by (Paudel et al., 2024a). Onion seeds (*Allium cepa* var. Texas Grano 1015Y) were surface sown on Sungrow 3B potting mix in 3-inch pots. After covering seeds with a layer of potting soil, pots were maintained in growth chamber conditions at ∼27^0^C, 40-50% humidity and watered 1-2 times per week as needed.

### Construction of *Burkholderia* and *Pantoea altR* promoter reporter constructs

The promoter capture vector pTn5.*7*luxk6 (GenBank: KC332287.1) was utilized to design *altR* promoter reporter construct for Bga strain 20GA0385 and *P. ananatis* strain PNA 97-1R following the procedure described by Jan et al., 2025 (Jan et al., 2025). The 77-bp sequence region containing the predicted *altR* promoter of PNA97-1R, along with an additional 30 bp upstream sequence of the StuI site and 30 bp downstream sequence of the DraIII site in the plasmid, was synthesized as double-stranded DNA (dsDNA) by Twist Biosciences. For the Bga *altR* reporter construct, 237 bp sequence region of the predicted 20GA0385 *altR* promoter region was amplified using primer pair pburkaltR-F and pburkaltR-R. The synthesized double-stranded DNA and PCR-amplified gel-extracted product were stitched to gel purified StuI and DraIII double-digested vector using Gibson assembly master mix (NEB, Ipswich MA) following manufacturer’s recommendations. The Gibson reaction product was transformed into electrocompetent MahI cells and selected on LB agar plates amended with kanamycin. The bioluminescent clones containing the *P. ananatis* P*altR* insert were plasmid purified using GeneJet Plasmid Miniprep kit (Thermoscientific, Watham, WA) and sequence confirmed using Sanger sequencing from Eurofins Genomics (Louisville, KY). The plasmid containing Bga P*altR* promoter construct was first confirmed by PCR using primer pair Tn*7*F and pBurkaltR-R. The representative clones that gave a PCR amplicon size of around 250 bp were purified using Monarch NEB PCR cleanup kit (New England Biolabs, Ipswich, MA) and confirmed using sanger sequencing from Eurofins Genomics and by using whole plasmid sequencing from Plasmidsaurus (Plasmidsaurus, Eugene, OR). The sequence confirmed plasmid was transformed into electro competent RHO5 cells. RHO5 donors with *altR* reporter plasmid construct and RHO3/pTNS3 transposition helpers were conjugated with recipient Bga 20GA0385 and PNA 97-1R strains and selected on LB+Rf+Kan plates. *P. ananatis* transposition candidates were screened for the insert using zone of inhibition assays. Bga transposition candidates were screened for the insert by zone of inhibition assays and PCR using primer pairs glmSR/Tn7LF and glmSR/pburkaltR-R with expected product size of 356 bp and 657 bp respectively.

### Construction of a constitutive *lux* reporter construct

The *ompA* promoter capture vector pTn7oLuxK4 (GenBank: KC332284.1) was used to build constitutively bioluminescent Bga 20GA0385 WT and ΔTTG strains (Bruckbauer et al., 2015). The P*_ompA_* based construct was utilized as a thiosulfinate-independent metabolic signal to monitor bacterial viability *in planta* independently of P*_altR_*de-repression. The plasmid harbored in MaH1 cells was purified and transformed into electrocompetent RHO5 cells. The RHO5 donor with the plasmid and RHO3/pTns3 transposition helper was conjugated into recipient WT or TTG strains, and candidate clones were selected on LB+Rf+Kan plates. The candidate clones were screened for bioluminescence using Syngene Pxi imager (Discovery Scientific Solutions, AZ) under 2 minutes exposure.

### Preparation of the pantaphos cell-free spent medium

Pantaphos cell-free spent medium prepared from the IPTG-induced PNA97-1R P_SNARE_ *hvr* overnight cultures was used as a non-native necrosis-inducing factor in the onion red scale necrosis assays. The cell-free spent medium was prepared following the procedures described previously (Shin et al., 2023). Briefly, overnight cultures of IPTG-inducible WT strain PNA97-1R::P_SNARE_*hvr* and IPTG-inducible *hvrE* deletion mutant strain PNA97-1R Δ*hvrE* :: P_SNARE_ *hvr* were started in LM broth. The next day, 50 µl of overnight culture was subcultured into a new 5 ml of LM media and grown until the OD_600_ value reached 0.6. This was followed by the addition of 50 µl of 1mM IPTG and cultures were left to grow overnight with shaking. The next day, the cultures were centrifuged at 4^0^C for 10 min at 2,585 relative central force (Eppendorf 5810R). The supernatant was then filter-sterilized through a 0.2 µm filter. The filter-sterilized medium was then stored at 4^0^C prior to experimental use.

### Onion leaf infection assay

The de-repression of *Burkholderia altR* promoter reporter construct *in planta* was monitored across three days post-inoculation using an onion seedling assay. Onion seedlings aged 8-12 weeks were inoculated with 20GA0385 Tn*7*P*_altR_*^PNA 97-1^LuxK6 and constitutively bioluminescent 20GA0385 P*_ompA_*Tn*7*oLuxK4 construct 4, 3, 2, and 1 day before imaging. At least two leaves per seedling and a total of 3 seedlings were inoculated for each day before imaging. Three independent experimental repeats were performed. The bioluminescence in the inoculated seedling samples was visualized and analyzed using the Newton 7.0 in vivo optical imaging system (Scintica Instrumentation Inc, Webster, TX). Samples were cut and aligned in the imaging plane and incubated inside the machine in the dark for 5 minutes to minimize autofluorescence signal from green tissue. The samples were imaged using the program-recommended exposure settings. The images were exported from the machine software and then imported into Kuant v 4.5 software for further analysis. The photon count scale bar in the software was manually adjusted to ensure no background noise was observed in the seedlings. After adjusting the scale, the samples that showed bioluminescence signal were counted as positive. Samples that showed a bioluminescence signal over the total number of inoculated samples were reported for each day.

### Preparation of bacterial inoculum

Normalized bacterial suspension for the scale necrosis assay, scale assay, and foliar assay was prepared following the procedures described by Paudel et al., 2024a; 2024b (Paudel et al., 2024a; Paudel et al., 2024b). Briefly, strains were streaked on LB plates amended with appropriate antibiotics and incubated at 30^0^C for 24 hours. The streaked area was suspended with 300 µl of sterile H_2_0 and spread to produce bacterial lawns. The plate was further incubated for 24 h and the next day, the lawn was scraped and suspended in 1 ml of sterile H_2_0 and standardized to OD_600_ = 0.7 (∼6 x 10^8^ CFU/ml). The standardized suspension was used for inoculation.

### Onion scale necrosis assay

Onion red scale necrosis (RSN) assay was performed following the procedure described by (Paudel et al., 2024a; Paudel et al., 2024b). Consumer-produced red onions were cut into 3-5 cm x 3-5 cm pieces, soaked in 3 % sodium hypochlorite solution for 2 minutes, and rinsed in dH_2_0 for 5-6 times. Washed scales were dried on a paper towel and kept on top of a 200 µl pipette tip rack placed on top of ethanol-sterilized flats. A 200 µl sterile pipette tip was used to wound the scales in the center, and a normalized bacterial suspension of OD_600_ = 0.7, 10 µl was deposited into the wound. The inoculated scales were then covered and incubated at room temperature for 3 days.

For the red scale necrosis assays, 10 µl of cell free spent medium of PNA97-1R::P_SNARE_ *hvr* / PNA97-1R Δ*hvrE* ::P_SNARE_ *hvr* was mixed with normalized suspension of Bga WT Tn7P ^PNA 97-1R^LuxK6 and 20 µl of the mix was inoculated into scales. To monitor the colonization of Bga 20GA0385 strain in scale tissue in the presence of cell free spent medium, constitutively bioluminescent version of WT strain was mixed with cell free spent medium of PNA97-1R:: P_SNARE_ *hvr* / PNA97-1R Δ*hvrE* :: P_SNARE_ *hvr* and inoculated into wounded scales. A total of six scales were inoculated per strain and bioluminescence/de-repression in the scale was monitored from day 1 to day 3 post inoculation in 24 hours interval using analytikJena UV chemstudio imager. Exposure time of 2 minutes was used for samples 24 h post-inoculation, whereas for the rest of the days, 5 minutes exposure setting was used. Scales that showed visible bioluminescence were counted as positive. The total number of scales that showed the presence of bioluminescence over the total number of inoculated scales was reported for each day. A total of three independent experimental repeats were performed.

Bga 20GA0385 P*_ompA_*Tn*7*oLuxK4 and 20GA0385 ΔTTG P*_ompA_*Tn*7*oLuxK4 strains were mixed with cell-free medium as described above and inoculated into red scale tissue to count the recovery of Bga strains 24 to 72 hours post-inoculation. A total of six scales were inoculated per treatment and each day; four symptomatic scales were randomly processed for *in planta* population count. The samples were processed following the procedure as described by (Paudel et al., 2024a; Paudel et al., 2024b). Symptomatic area around the point of inoculation was sampled with an ethanol sterilized cork bore (*r* = 2.5 mm) and mixed with 200 µl sterile milliQ H_2_0 in a 2-ml SARSTEDT microtube (SARSTEDT AG & Co., Numbrecht, Germany) containing three 3-mm high density zirconium beads (Glen Mills grinding media). The sample was crushed in a Bead Ruptor Elite Bead Mill Homogenizer (Omni International, Kennesaw, GA) for 30 seconds at 4 m/s speed settings. The sample was serially diluted (20 µl, 180 µl) in a 96-well styrene plates and diluents were plated on LM media amended with rifampicin. The colony forming unit (CFUs) were back-calculated to determine CFUs per mg of infected sample. The experiment was repeated at least three times. The differences in population recovery between the WT and TTG mutant from the co-inoculation mix for each day were analyzed for significance using pairwise t-test function in RStudio v 2023.09.0.

### Preparation of onion juice

Onion juice was extracted from consumer produce yellow onions following methods reported previously (Stice et al., 2020). In brief, an onion bulb was processed using a consumer-grade juicer (Breville Juice Fountain Elite), producing approximately 200 to 300 mL of crude onion extract. The extract was then centrifuged at 14,000 × g for 2 hours at 4°C using a Sorvall RC5B Plus centrifuge (Marshall Scientific, Hampton, NH) in a 250-mL centrifuge bottle. The supernatant was sterilized through a Nalgene disposable 0.2-micron vacuum filter unit. The filtered juice was then aliquoted and stored at –20°C for up to one week before use in experiments.

### Onion scale preconditioning assay

Six yellow onion scales were inoculated with normalized suspensions of WT and TTG strains and incubated in room temperature for 3 days. The necrotized onion tissue around the point of inoculation with conditioned WT or TTG strain was sampled using ethanol sterilized cork bore (r = 2.5 mm) and mixed with 200 µl sterile water in a 2-ml SARSTEDT and crushed with homogenizer as described above. A volume of 88 µl of crushed sample from each scale was mixed with half strength onion juice to make a total volume of 5280 µl for one strain. Each well in the 100-well honeycomb plates was inoculated with 400 µl of the mix. Each strain was inoculated into a total of 12 wells. Six wells were inoculated with 200 µg/ml kanamycin as a normalization control. The absorbance values recorded for kanamycin treated wells were averaged and subtracted from each of the remaining six wells for each time point. For the non-conditioned controls, normalized bacterial suspension of WT and TTG mutant strain was mixed with half strength onion juice and 400 µl of the mix was loaded into each well for a total of six wells per strain. At least three wells were inoculated with half strength onion juice as the negative control. Average reading of negative control was subtracted from the individual reading of each well. The obtained absorbance readings across a 48-hour time period were plotted in a line graph using the ggplot function in Rstudio.

### Preparation of allicin stock solution

The allicin solution for the zone of inhibition assay was prepared as described by (Stice et al., 2020; Paudel et al., 2024b). Briefly, 25 µl of glacial acetic acid (Sigma Aldrich), 15 µl of diallyl disulfide 96% (Carbosynth), and 15 µl of 30% H_2_O_2_ were all mixed together in a 200 µl PCR tube. The PCR tube was sealed and agitated at 30^0^C for 6 h. After 6 h, the tube was centrifuged for 10 seconds and suspended in 1 ml of methanol to halt the reaction. The mix was used directly as a synthesized allicin stock for the zone of inhibition assay. A fresh stock was made for each experiment.

### Zone of inhibition assays

The de-repression of Bga P*_altR_*^PNA 97-1R^Tn*7*LuxK6 and Bga P*_altR_*^20GA0385^Tn*7*LuxK6 strains was tested using allicin ZOI assay. 5 mL overnight cultures were started from a single colony in LBKan broth. 300 µl of the culture was spread in LBKan plates, and synthesized allicin was added in the hole punched at the center of the agar plate. The plates were incubated for 18 h and visualized for bioluminescence using analytikJena imager under 2-or 5-minute exposure. This experiment was repeated twice.

To check whether the Bga 20GA0385 WT strain can rescue the *in vitro* phenotype of its TTG mutant derivative, a ZOI co-inoculation assay was done. Overnight cultures (5 ml, ∼24 h) of WT or ΔTTG P*_ompA_*Tn*7*oLuxK4 strain were started in LB media amended with rifampicin and equal volume of both strains was mixed. For WT only and ΔTTG only treatment, cultures were half diluted with sterile LB. The suspension of 400 µl volume was spread to make a lawn on LB agar and left to dry for 10 minutes. Agar hole was punched in the center of the plate with the back end of a sterile 20 µl Rainin pipette tips. The hole was then inoculated with 50 µl of synthesized allicin and the plates were incubated in 30^0^C for 24 hours. Next day, brightfield and bioluminescence images of the plates were visualized using analytiKjena imager. Exposure settings of 2 min was used for bioluminescence. The inhibition area was measured using ImageJ 1.54g software. The difference in inhibition area between the treatments was analyzed for level of significance using pairwise t-test function in R studio. The ZOI area between different strains was presented in a box plot generated using ggplot function in R studio.

### Onion seedling and leaf co-inoculation assay

The co-inoculation assay was done with 8-12 weeks old seedling to test if the Bga 20GA0385 strain can rescue the population of ΔTTG strain *in planta*. Seedlings were marked approximately at the midpoint for inoculation and 0.5 cm up and down of the inoculation point for processing. Normalized bacterial suspension for the WT and ΔTTG Tn*7*oLuxK4 strain was prepared as described in the above section. For co-inoculation mix, equal volume of normalized bacterial suspension was mixed and 20 µl of the mix was inoculated into the seedling. For WT and TTG standalone treatments, normalized suspension was half diluted with sterile H_2_0 and 20 µl of the mix was inoculated. After 3 days post inoculation, necrosis length was measured, and samples were excised from the pre-marked area and processed for CFUs count following the procedure as described above for the onion scale necrosis assay. The CFU recovery was normalized per cm length of the sampled tissue. The CFU of TTG mutant from the coinoculation mix was determined by counting the bioluminescent clones from the serial diluted co-inoculated tissue macerate suspension plated on LM agar amended with rifampicin. One seedling leaf per plant per pot and a total of six pots were inoculated per treatment. The experiment was repeated three times in its entirety.

Mature onion plants were also inoculated to test if the Bga WT strain can rescue the TTG mutant population in the co-inoculation mix. Small onion sets (cv. Century) were transferred to 4-inch pots and allowed to grow for 9-11 weeks under greenhouse conditions before inoculation. Two oldest leaves per plant were inoculated and a total of 3 plants were inoculated per treatment. The leaf samples were marked and processed for CFU count as described above. The CFU/mg reading of two leaves per plant was averaged and used as a single data point for further analysis. The experiment was repeated six times in its entirety.

## ACKNOWLEDGMENTS

This manuscript was edited with assistance from Microsoft Co-Pilot.

## Supplemental Figures

**Fig. S1**. The derepression of Bga *altR* reporter construct is seen as a bright ring in the ZOI assay. Representative brightfield and bioluminescence images of **Top**, Bga P*_altR_*^PNA 97-1R^Tn*7*LuxK6 strain and **Bottom**, Bga P*_altR_*^20GA0385^Tn*7*LuxK6. Images taken 18 hours post inoculation. Scale: 1 cm.

**Fig. S2**: Bga WT strain didn’t rescue the population of TTG mutant from the infected mature onion plants in the co-infection assay. Box plot showing the CFU/mg recovery of Bga WT, TTG mutant, and TTG mutant from co-infection mix after three days post inoculation. n is the total number of observations across six independent experimental repeats.

## REFERENCES

Belo, T., du Toit, L., Waters, T., Derie, M., Schacht, B., and LaHue, G. 2023. Reducing the risk of onion bacterial diseases through managing irrigation frequency and final irrigation timing. Agricultural Water Management 288:108476.

Borlinghaus, J., Bolger, A., Schier, C., Vogel, A., Usadel, B., Gruhlke, M., and Slusarenko, A. 2020. Genetic and molecular characterization of multicomponent resistance of *Pseudomonas* against allicin. Life science alliance 3.

Bruckbauer, S. T., Kvitko, B. H., Karkhoff-Schweizer, R. R., and Schweizer, H. P. 2015. Tn5/7-lux: a versatile tool for the identification and capture of promoters in gram-negative bacteria. BMC Microbiol. 15:17.

Cho, H., Park, J., Kim, D., Han, J., Natesan, K., Choi, M., Lee, S., Kim, J., Cho, K., and Ahn, B. 2024. Understanding the defense mechanism of *Allium* plants through the onion isoallicin-omics study. Frontiers in Plant Science 15:1488553.

Eady, C.C., Kamoi, T., Kato, M., Porter, N.G., Davis, S., Shaw, M., Kamoi, A., and Imai, S. 2008. Silencing onion lachrymatory factor synthase causes a significant change in the sulfur secondary metabolite profile. Plant physiology 147:2096–2106.

Hughes, J., Tregova, A., Tomsett, A., Jones, M., Cosstick, R., and Collin, H. 2005. Synthesis of the flavour precursor, alliin, in garlic tissue cultures. Phytochemistry 66:187–194.

Jan, H., Kong, F., MacLellan, M.P., Yang, L., Dutta, B., and Kvitko, B. 2025. The AltR transcription factor responds to plant thiosulfinates to regulate gene expression in a bacterial pathogen of onion. bioRxiv:2025.2011.2019.689302.

Kvitko, B.H., Bruckbauer, S., Prucha, J., McMillan, I., Breland, E.J., Lehman, S., Mladinich, K., Choi, K., Karkhoff-Schweizer, R., and Schweizer, H. 2012. A simple method for construction of pir+ Enterobacterial hosts for maintenance of R6K replicon plasmids. BMC research notes 5:1–7.

Lancaster, J.E., and Collin, H.A. 1981. Presence of alliinase in isolated vacuoles and of alkyl cysteine sulphoxides in the cytoplasm of bulbs of onion (*Allium cepa*). Plant Science Letters 22:169–176.

Lee, C.J., Lee, J.T., Kwor, J., Kim, B., and Park, W. 2005. Occurrence of bacterial soft rot of onion plants caused by *Burkholderia gladioli* pv. *alliicola* in Korea. Australasian Plant Pathology 34:287–292.

Leontiev, R., Hohaus, N., Jacob, C., Gruhlke, M., and Slusarenko, A.J. 2018. A comparison of the antibacterial and antifungal activities of thiosulfinate analogues of allicin. Scientific reports 8:6763.

López, C.M., Rholl, D.A., Trunck, L.A., and Schweizer, H.P. 2009. Versatile dual-technology system for markerless allele replacement in *Burkholderia pseudomallei*. Applied and environmental microbiology 75:6496–6503.

Müller, A., Eller, J., Albrecht, F., Prochnow, P., Kuhlmann, K., Bandow, J.E.S., Alan, J., and Leichert, L. 2016. Allicin induces thiol stress in bacteria through S-allylmercapto modification of protein cysteines. Journal of Biological Chemistry 291:11477–11490.

Paudel, S., Franco, Y., Zhao, M., Dutta, B., and Kvitko, B.H. 2024a. Distinct Virulence Mechanisms of *Burkholderia gladioli* in Onion Foliar and Bulb Scale Tissues. Molecular Plant-Microbe Interactions.

Paudel, S., Zhao, M., Stice, S.P., Dutta, B., and Kvitko, B.H. 2024b. Thiosulfinate Tolerance Gene Clusters Are Common Features of *Burkholderia* Onion Pathogens. Mol Plant Microbe Interact 37:507–519.

Reiter, J., Hübbers, A., Albrecht, F., Leichert, L., and Slusarenko, A. 2020. Allicin, a natural antimicrobial defence substance from garlic, inhibits DNA gyrase activity in bacteria. International Journal of Medical Microbiology 310:151359.

Rose, P., Whiteman, M., Moore, P.K., and Zhu, Y. 2005. Bioactive S-alk (en) yl cysteine sulfoxide metabolites in the genus *Allium*: the chemistry of potential therapeutic agents. Natural product reports 22:351–368.

Schwartz, H., du Toit, L., and Coutinho, T. 2015. Diseases of Onion and Garlic (*Allium cepa* L. and *A. sativum* L., Respectively). The American Phytopathological Society: St Paul, MN, USA.

Shin, G., Dutta, B., and Kvitko, B.H. 2023. The Genetic Requirements for HiVir-Mediated Onion Necrosis by *Pantoea ananatis*, a Necrotrophic Plant Pathogen. Molecular Plant-Microbe Interactions® 36:381–391.

Stice, S., Thao, K., Khang, C., Baltrus, D., Dutta, B., and Kvitko, B. 2020. Thiosulfinate tolerance is a virulence strategy of an atypical bacterial pathogen of onion. Current Biology 30:3130–3140. e3136.

Van Damme, E., Smeets, K., Torrekens, S., Van Leuven, F., and Peumans, W.J. 1992. Isolation and characterization of alliinase cDNA clones from garlic (*Allium sativum* L.) and related species. European Journal of Biochemistry 209:751–757.

Yamazaki, Y., Iwasaki, K., Mikami, M., and Yagihashi, A. 2010. Distribution of eleven flavor precursors, S-alk (en) yl-L-cysteine derivatives, in seven *Allium* vegetables. Food science and technology research 17:55–62.

