## Supplementary figures and images for "Evidence for *Burkholderia gladioli* pv. *alliicola* Extracellular Detoxification of Thiosulfinates"

### Fig. S1

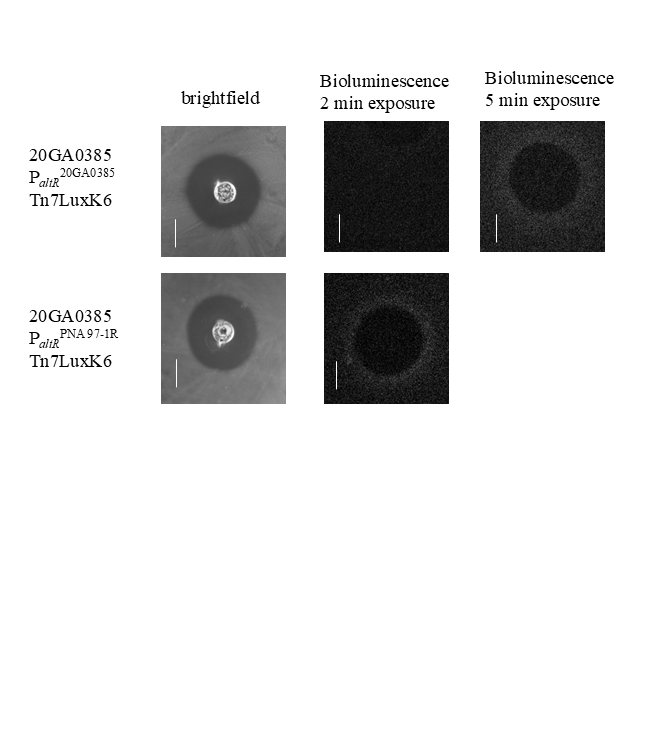

### Fig. S2

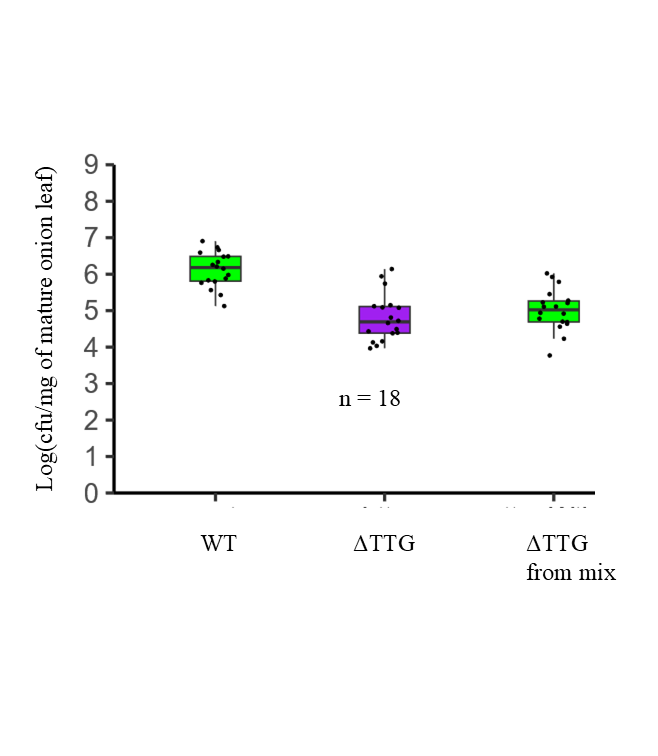
